# TRIP6 inhibits the Hippo signaling pathway in response to tension at adherens junctions

**DOI:** 10.1101/182204

**Authors:** Shubham Dutta, Sebastian Mana-Capelli, Murugan Paramasivam, Ishani Dasgupta, Heather Cirka, Kris Billiar, Dannel McCollum

**Author notes:** Address correspondence to: Dannel McCollum, University of Massachusetts Medical School, 364 Plantation Street, Worcester, MA, 01605.

## Abstract

The transcriptional co-activator YAP controls cell proliferation, survival, and tissue regeneration in response to changes in the mechanical environment. It is not known how mechanical stimuli such as tension are sensed and how the signal is transduced to control YAP activity. Here we show that the LIM domain protein TRIP6 acts as part of a mechanotransduction pathway at adherens junctions to promote YAP activity by inhibiting the LATS1/2 kinases. Previous studies showed that vinculin at adherens junctions becomes activated by mechanical tension. We show that vinculin inhibits Hippo signaling by recruiting TRIP6 to adherens junctions and stimulating its binding to and inhibition of LATS1/2 in response to tension. TRIP6 competes with MOB1 for binding to LATS1/2 thereby blocking MOB1 from recruiting the LATS1/2 activating kinases MST1/2. Together these findings reveal a novel mechanotransduction cascade that transduces tension signals sensed at adherens junctions to control Hippo pathway signaling.

## Introduction

Tissue architecture and mechanical forces are major regulators of cell proliferation, and they play important roles during development, organ growth, and tissue regeneration (Heller and Fuchs, 2015; Huang and Ingber, 1999; Mammoto et al., 2013). The cytoskeleton, extracellular matrix, and cell-cell adhesion are critical for transmitting force between cells and across tissues (Vogel and Sheetz, 2006). The Hippo signaling pathway is a major regulator of cellular responses to mechanical inputs (Halder et al., 2012; Sun and Irvine, 2016). The core Hippo pathway (Meng et al., 2016) consists of two kinase modules: the first includes several Ste20-superfamily kinases (MST1/2 are the best characterized), which phosphorylate and activate the LATS1/2 kinases. MST1/2 phosphorylation of LATS1/2 is mediated by MOB1, which promotes association of MST1/2 with LATS1/2. LATS1/2 then phosphorylate and inhibit the transcriptional co-activator YAP (and its homolog TAZ) by causing it to be sequestered in the cytoplasm or degraded. When in the nucleus, YAP associates with transcription factor TEAD to upregulate genes responsible for survival, proliferation, and stem cell maintenance. The growth promoting properties of YAP are frequently co-opted by cancer cells, in which YAP is often activated and overexpressed (Yu et al., 2015). Although the activity of both LATS1/2 and YAP are clearly regulated by mechanical inputs, how those inputs are sensed and the signals are transduced remain obscure.

Experiments in *Drosophila* and mammalian cells revealed that Hippo pathway regulation of YAP is controlled by mechanical tension (Aragona et al., 2013; Benham-Pyle et al., 2015; Codelia et al., 2014; Rauskolb et al., 2014). When cells experience high mechanical tension, YAP localizes to the nucleus and promotes cell proliferation. Conversely, low tension causes YAP to exit the nucleus and cells to arrest growth. Transmission of tension across tissues requires cell-cell adhesion such as that provided by cadherins (Mui et al., 2016). Tension experienced by cells can be generated by the cells themselves through actomyosin stress fibers or by externally imposed stretch or force (Halder et al., 2012). Studies in *Drosophila* indicate that tension within tissues decreases as cell density increases, and hence tension sensing could contribute to density dependent inhibition of cell growth, a property that is typically lost in cancer cells (Rauskolb et al., 2014). Perturbation of stress fibers, externally applied stretch, and cell density all modulate LATS1/2 and YAP activity; however, the sensors are not known. In *Drosophila,* the LIM domain protein Ajuba inhibits Warts (the LATS1/2 homolog) and recruits it to adherens junctions in a tension-dependent manner (Rauskolb et al., 2014). The mechanism by which Ajuba regulates Warts activity is not clearly understood, and reports have differed regarding whether Ajuba-related proteins function similarly in mammals (Codelia et al., 2014; Jagannathan et al., 2016). Here we show that the human LIM domain protein TRIP6 acts as part of a mechanotransduction cascade at adherens junctions to regulate LATS1/2 in response to mechanical tension at cell-cell junctions.

## RESULTS

### TRIP6 activates YAP through inhibition of LATS1/2

Although TRIP6 is overexpressed in various cancers where it promotes proliferation and invasion (Chastre et al., 2009; Fei et al., 2013; Grunewald et al., 2013), prior studies had not connected TRIP6 to the Hippo signaling pathway. We previously identified TRIP6 as one of several LATS2 binding partners using tandem affinity purification and mass spectrometry (Paramasivam et al., 2011). Here, to validate the LATS2-TRIP6 interaction, we performed co-immunoprecipitation experiments. LATS2 was pulled down in TRIP6 immunoprecipitates when both proteins were overexpressed (Figure 1A). In addition, endogenous LATS1 was present in TRIP6 immune complexes isolated from MCF10A cells (Figure 1B). Like its related family members (Zyxin, LPP, Ajuba, WTIP, and LIMD1), the carboxy-terminal half of TRIP6 consists of 3 conserved LIM domains (Figure 1A). Truncation experiments showed that LATS2 binding maps to the C-terminal LIM domain half of TRIP6 (Figure 1A). We next tested which parts of LATS2 interacted with TRIP6. TRIP6 bound to the N-terminal region of LATS2 and specifically interacted with two segments (amino acids 376-397 and 625-644) (Figure 1C). This finding is highly reminiscent of previous findings with Ajuba and Zyxin (Abe et al., 2006; Hirota et al., 2000).

**Figure 1.**
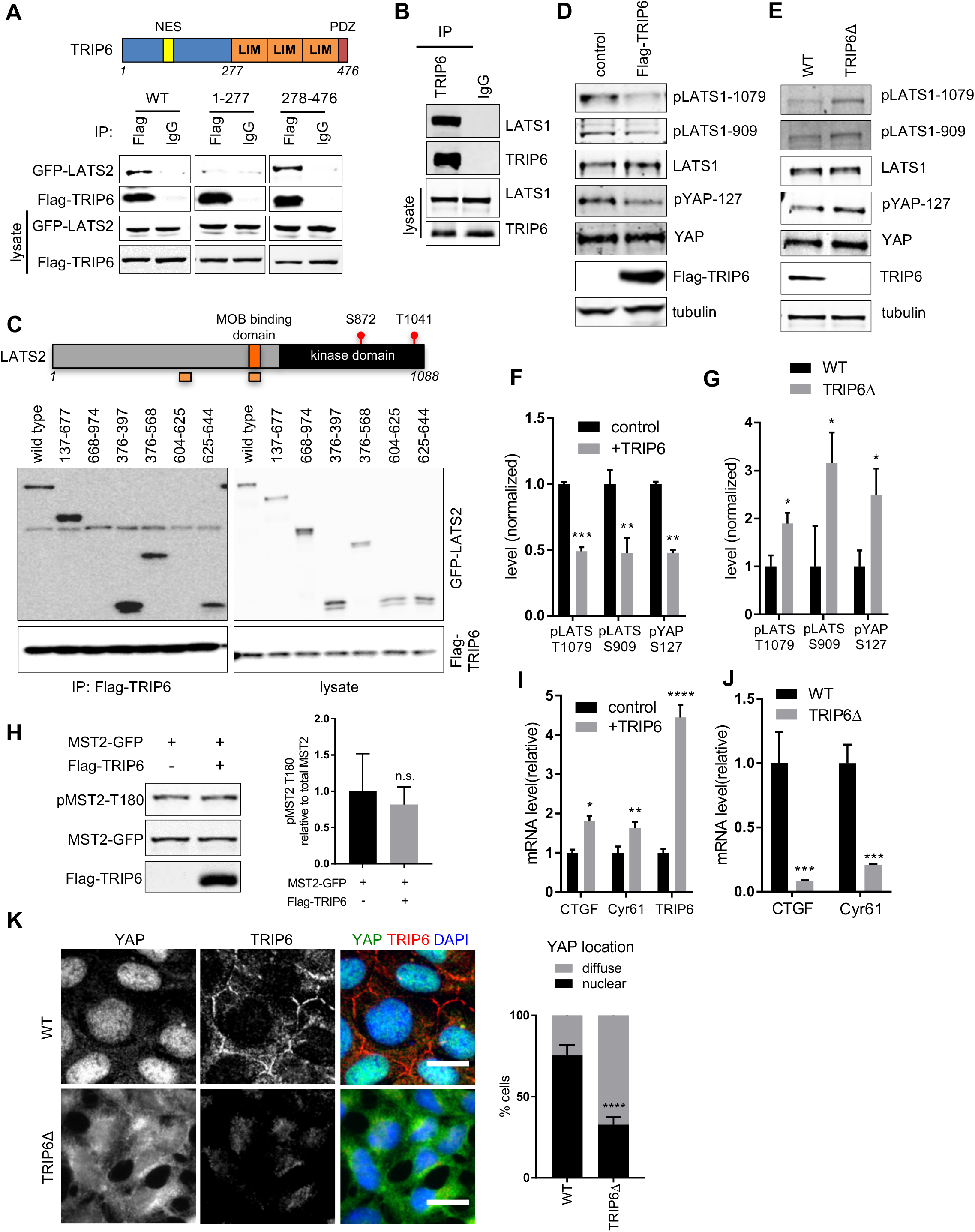
TRIP6 promotes YAP activity by inhibiting LATS1/2 (A) Full length (WT), the amino-terminal half (1-277), or the carboxy-terminal half (278-476) of TRIP6 were tested for binding to LATS2 by immunoprecipitation. FLAG-TRIP6 variants were co-expressed with LATS2-GFP in HEK293 cells, anti-FLAG or control (IgG) antibodies were used to isolate immune complexes. Immune complexes and lysates were probed by western blotting for LATS-GFP and FLAG-TRIP6. Schematic diagram depicts TRIP6 domains (NES: Nuclear Export Signal). (B) Lysates from MCF10A cells were subjected to immunoprecipitation using anti-TRIP6 or control (IgG) antibodies, and immune complexes and lysates were probed for TRIP6 and LATS1. (C) FLAG-TRIP6 was tested for binding to various LATS2-GFP deletion mutants as described in part (A). Schematic diagram of LATS2 shows MOB1 binding domain, and the autophosphorylation (S872) and MST1/2 phosphorylation sites (T1041) in the kinase domain. The regions marked in green depict TRIP6 binding sites on LATS2 (D) Lysates from HEK293A cells transfected with control or FLAG-TRIP6 plasmid were analyzed by western blotting using the indicated antibodies (quantification is shown in (F)). (E) Lysates from control (WT) or CRISPR generated TRIP6 null (TRIP6A) HEK293A cells were analyzed by western blotting using the indicated antibodies (quantification shown in (G)). (F) The relative levels of LATS1 activating phosphorylation (pLATS1-1079, 909) and YAP S127 inhibitory phosphorylation from (D) were measured relative to LATS1 and YAP levels respectively. (Mean ± SD; n=3; **P<0.01, ***P<0.001, T-test) (G) The levels of LATS1 activating phosphorylation and YAP inactivating phosphorylation from part (E) were quantified (Mean ± SD; n=3; *P<0.05, T-test). (H) GFP-MST2 was expressed with or without FLAG-TRIP6 in HEK293 cells and the levels of MST2, MST2 activating phosphorylation (pMST2-T180), and FLAG-TRIP6 were measured by western blotting with the indicated antibodies. (Mean ± SD; n=3; n.s.>0.05, T-test). (I) TRIP6 was overexpressed in HEK293A cells and the levels of TRIP6, and YAP target gene expression was analyzed using RT-qPCR. (Mean ± SD; n=3; *P<0.05, **P<0.01, ****P<0.0001, T-test). (J) The levels of YAP target gene expression was analyzed using RT-qPCR in control (WT) and TRIP6A HEK293A cells. (Mean ± SD; n=3; ***P<0.001, T-test) (K) Control (WT) and TRIP6A HEK293A cells were stained for YAP and TRIP6. Merged image shows YAP (green), TRIP6 (red), and DNA (blue). Quantification of at least 100 cells is shown (Mean ± SD; n=3; ****P<0.0001, Fisher’s test). Scale bar=20µm.

To determine whether TRIP6 regulates LATS1/2 activity, we examined the effects of TRIP6 overproduction and loss of function. Overproduction of TRIP6 in HEK293A cells reduced endogenous LATS1/2 activity as judged by probing the two sites of activating phosphorylation on LATS1/2, T1079 and S909 (for LATS1, T1041 and S872 for LATS2) (Figure 1D & F) (note that T1079 is phosphorylated by MST1/2 and S909 is an autophosphorylation site). In contrast, TRIP6 overexpression did not affect MST2 activating phosphorylation (Figure 1H), suggesting that TRIP6 may regulate the ability of LATS1/2 to be phosphorylated by MST1/2. CRISPR mediated deletion of TRIP6 in HEK293A cells (Figure 1E & G) or shRNA mediated knockdown of TRIP6 in MCF10A cells (Figure S1A) increased LATS1/2 activating phosphorylation levels. Together these results show that TRIP6 acts to inhibit LATS1/2 activity.

Because LATS1/2 phosphorylate and inhibit YAP nuclear localization, stability, and activity, we tested the effect of modulating TRIP6 levels on YAP. Overexpression of TRIP6 in HEK293A cells inhibited LATS1/2 phosphorylation of YAP on S127 (Figure 1D & F) and increased expression of YAP target genes (Figure 1I). In contrast, reduced levels of TRIP6 inhibited YAP function. Specifically, shRNA mediated knockdown of TRIP6 in MCF10A cells reduced expression of YAP target genes (Figure S1B), and diminished YAP nuclear localization (Figure S1C). These cells also had reduced levels of YAP protein (Figure S1D), presumably caused by LATS1/2 phosphorylation dependent degradation (Liu et al., 2010; Zhao et al., 2010). TRIP6Δ HEK293A cells showed increased YAP S127 phosphorylation (Figure 1E & G), reduced expression of YAP target genes (Figure 1J), and reduced YAP nuclear localization (Figure 1K). Our observation that MCF10A, but not HEK293A, cells had reduced levels of YAP when TRIP6 was depleted (or eliminated) may reflect cell type differences in YAP degradation in response to LATS1/2 dependent phosphorylation. The TRIP6Δ HEK293A cells also displayed a defect in cell-cell adhesion as judged by the presence of frequent gaps between cells even at high density that were not observed in parental HEK293A cells (Figure 1K, S1E). The cell-cell adhesion and YAP localization defect in TRIP6Δ HEK293A cells was rescued by reexpression of TRIP6 (Figure S1E-F). MCF10A cells knocked down for TRIP6 with shRNA did not show obvious cell-cell adhesion defects perhaps due to the presence of residual TRIP6. Overall, these results show that TRIP6 inhibition of LATS1/2 promotes YAP activity.

### TRIP6 inhibits LATS1/2 by blocking binding to MOB1

We next investigated the mechanism for how TRIP6 inhibits LATS1/2. TRIP6-related LIM domain proteins have been shown to bind and inhibit LATS (Abe et al., 2006; Das Thakur et al., 2010; Hirota et al., 2000; Rauskolb et al., 2011; Rauskolb et al., 2014; Reddy and Irvine, 2013), however, it is not clear how they regulate LATS1/2 activity. Although zyxin was shown to promote degradation of LATS1/2 in response to hypoxia (Jagannathan et al., 2016), we did not observe any changes in LATS1 levels when TRIP6 levels were altered suggesting that TRIP6 uses a different mechanism. Because one of the TRIP6 binding sites in LATS2 (amino acids 625-644) overlaps with the binding site for its activator MOB1 (amino acids 595-662)(Ni et al., 2015), we wondered if TRIP6 and MOB1 compete for binding to LATS1/2. This mechanism would be consistent with our observations that TRIP6 inhibits the ability of MST1/2 to phosphorylate LATS1/2, because MOB1 activates LATS1/2 by promoting its association with and phosphorylation by MST1/2 (Ni et al., 2015). We first examined if TRIP6 could inhibit LaTs1/2-MOB1 binding *in vivo.* We found that TRIP6 overexpression reduced LATS2-MOB1A association in HEK293 cells (Figure 2A). To determine whether TRIP6 directly competes with MOB1A for binding to LATS2, competition experiments were carried out using purified recombinant proteins. Initial results demonstrated that GST-TRIP6 bound directly to MBP-LATS2 but not MBP alone (Figure 2B, compare lanes 1 and 3). Competition experiments showed that MOB1A could compete with TRIP6 for binding to LATS2. 6HIS-MOB1A bound to MBP-LATS2 and inhibited GST-TRIP6 binding, with the highest levels of MOB1A reducing TRIP6-LATS2 binding to background levels (Figure 2B, lanes 3-6). Addition of non-specific competitor (BSA), at the same level as the highest amount of MOB1A used (Figure S2), did not cause any reduction in TRIP6-LATS2 binding (Figure 2B, lane 7). Together these results show that that TRIP6 and MOB1 compete for binding to LATS2 and that TRIP6 likely inhibits LATS1/2 activity at least in part by blocking MOB1 binding.

**Figure 2.**
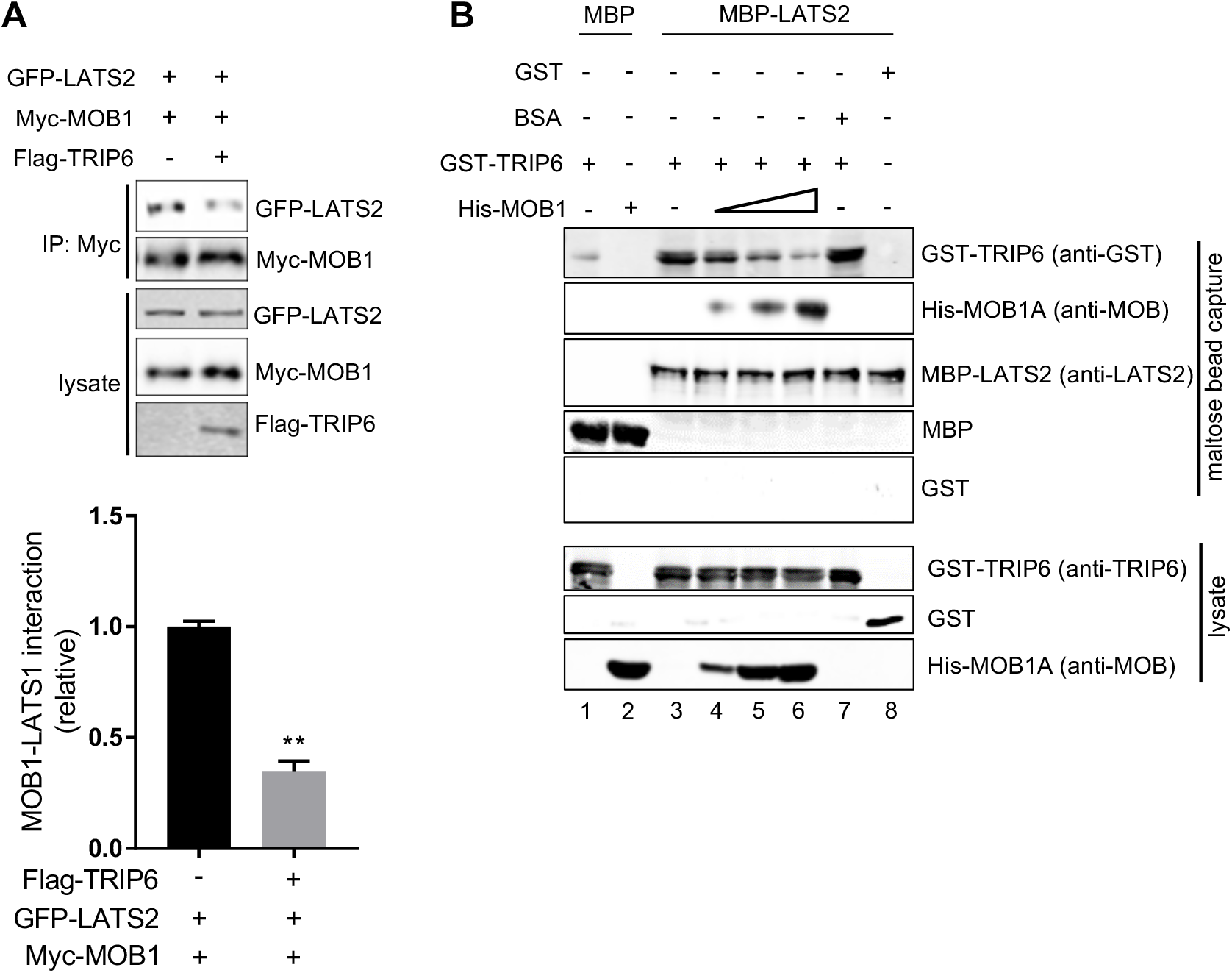
TRIP6 competes with MOB1 for binding to LATS2. (A) LATS2-GFP and MOB1A-Myc were overexpressed in HEK293 cells with or without co-overexpression of FLAG-TRIP6. MOB1A-Myc was immunoprecipitated using anti-Myc antibodies and assayed for MOB1A-Myc and LATS2-GFP levels. Levels of FLAG-TRIP6, MOB1A-Myc, and LATS2-GFP in the lysate are also shown. The levels of LATS2-GFP in immune complexes relative to the level of MOB1A-Myc are shown in the graph (Mean ± SD; n=3; **P<0.01, T-test) (B) Competitive binding experiments were done using purified recombinant MBP-LATS2, GST-TRIP6, and 6His-MOB1A. MBP-LATS2 bound to maltose beads was incubated with GST-TRIP6 with or without increasing amounts of 6His-MOB1A, and the levels of each protein bound to MBP-LATS2 on the beads at the end of the experiment was determined by western blotting. The levels of input proteins are shown (lysate). The binding of 6His-MOB1A and GST-TRIP6 to MBP alone, and the use BSA as a competitor instead of 6His-MOB1A are shown as controls. The numbers at the bottom are referred to in the text.

### TRIP6 modulates LATS1/2 activity and localization in response to tension at cell-cell junctions

To investigate what regulatory inputs might control TRIP6 inhibition of LATS1/2, we examined the localization of each protein. Endogenous TRIP6 and LATS1 co-localize to cell-cell junctions in MCF10A (Figure 3A, S3A) and to a lesser extent in HEK293A (Figure S3B) cells. Although we have been unable to find LATS2 antibodies capable of detecting the endogenous protein, GFP-LATS2 fusions also localize to cell-cell junctions (Paramasivam et al., 2011). TRIP6 has been previously reported to localize to both cell-cell junctions (adherens junctions) (Guo et al., 2014) and to focal adhesions (Wang et al., 1999; Zhao et al., 1999). Although we could faintly observe TRIP6 at focal adhesions in MCF10A cells at low density or at the edge of monolayers, at densities typically used in this study (confluent but still proliferating), TRIP6 was primarily at adherens junctions, and we saw little focal adhesion staining for TRIP6 or the focal adhesion marker FAK (Figure S3C). LATS1 was not observed at focal adhesions in MCF10A cells at any density (data not shown). We next assessed the mutual dependence of LATS1 and TRIP6 localization. Knockdown of TRIP6 in MCF10A cells (Figure 3A; S3D) reduced localization of LATS1 to cell junctions. Deletion of TRIP6 in HEK293A cells (Figure S3B) also caused reduced localization of LATS1 to cellcell junctions, although because of the reduced cell-cell adhesion in these cells, it is possible that effects on LATS1 localization could be due to defects in cell-cell junctions. When LATS1/2 were knocked down (depleted) in MCF10A cells, TRIP6 remained at cell-cell junctions (Figure 3B; S3E), but its localization was more punctate and less smooth, possibly reflecting a transition to a more mesenchymal state (Zhang et al., 2008). Together these results show that TRIP6 and LATS1/2 affect each other’s localization and that TRIP6 may recruit LATS1/2 to cell junctions.

**Figure 3.**
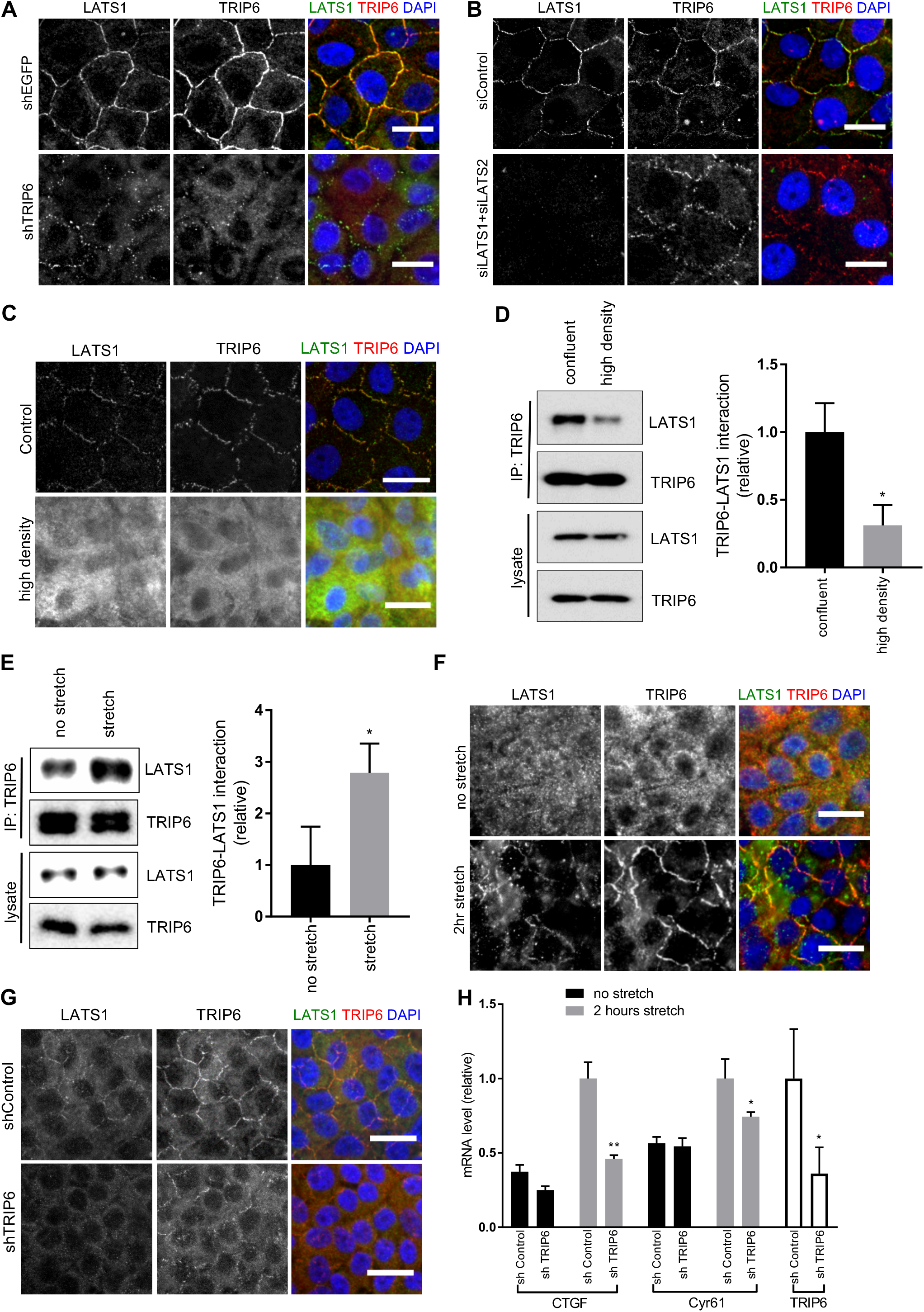
TRIP6-LATS binding and localization to cell-cell junctions is regulated by tension. (A) MCF10A cells were infected with lentivirus carrying control shRNA (shEGFP), or a mix of two different shRNA against TRIP6 (shTRIP6-1 and shTRIP6-4) and were stained for TRIP6 and LATS1. Merged images show LATS1 (green), TRIP6 (red) and DNA (blue) (quantification of LATS1 localization at cell-cell junctions is shown in Figure S3D). Scale bar=20µm. (B) LATS1 and LATS2 were knocked down using siLATS1 and siLATS2 SMARTPools in MCF10A cells. MCF10A control cells were treated with control siRNA (siControl) against fire fly luciferase. Cells were stained for TRIP6 and LATS1 as in (A) (quantification of TRIP6 localization at cell-cell junctions is shown in Figure S3E). Scale bar=20µm. (C) MCF10A cells were grown to high density, and were stained for TRIP6 and LATS1 as in (A). Scale bar=20µm. (D) Cells were grown as in (C), then lysed and anti-TRIP6 or control (IgG) antibodies were used to isolate immune complexes. Immune complexes and lysates were probed by western blotting for LATS1 and TRIP6. Quantification is shown. (Mean ± SD; n=3; *P<0.05, T-test). (E) MCF10A cells grown at high density on PDMS membranes and were stretched (or not) at 17% elongation for 2 hours and lysed while under tension. Anti-TRIP6 or control (IgG) antibodies were used to isolate immune complexes. Immune complexes and lysates were probed by western blotting for LATS1 and TRIP6. Quantification is shown. (Mean ± SD; n=3; *P<0.05, T-test). (F) Cells were treated as in (E), fixed while under tension, and stained for TRIP6 and LATS1 as in (A). Scale bar=20µm. (G) MCF10A cells infected with lentivirus carrying control shRNA (shEGFP), or a mix of two different shRNA against TRIP6 (shTRIP6-1 and shTRIP6-4), were grown at high density on PDMS membranes and were stretched or not (only stretched cells shown) at 17% elongation for 2 hours, fixed while under tension, and were stained for TR|P6 and LATS1 as in (A). Scale bar=20µm. (H) Cells were treated as in (G) and YAP target gene (CTGF and Cyr61) and TR|P6 expression were analyzed using RT-qPCR. (Mean ± SD; n=3; *P<0.05, **P<0.01, T-test).

We next examined whether recruitment of TRIP6 and LATS1 to cell junctions is regulated by stimuli that control LATS1/2 activity. Both TRIP6 and LATS1 localized to cell-cell junctions in cells that were confluent but still proliferating. However, in highly dense non-proliferating cells TRIP6 and LATS1 no longer localized to cell-cell junctions (Figure 3C, S3B), despite unchanged levels of both proteins (Figure 3D). Interestingly we also observed a reduction in TRIP6-LATS1 binding in MCF10A cells at high cell density (Figure 3D), consistent with the increased LATS1/2 activity observed under these conditions (Meng et al., 2015). How cell density controls TrIP6-LATS1/2 binding and localization is not clear. However, a study in *Drosophila* tissue showed that tension at cell-cell junctions is reduced as cell density increases (Rauskolb et al., 2014). Therefore, we tested whether increasing tension at cell-cell junctions in dense cultures would restore localization of TRIP6 and LATS1 to cell-cell junctions. To do this, we examined TRIP6 and LATS1 localization in dense cultures grown on flexible PDMS substrates before and after static stretch for 2 hours. We observed that stretch increased TRIP6-LATS1 binding (Figure 3E), localization of both proteins to cellcell junctions (Figure 3F), and YAP activity (Figure S3F). Both tension dependent recruitment of LATS1 to cell-cell junctions and YAP activation in dense monolayers following stretch depended on TRIP6 (Figure 3G-H). Together these results show that tension can trigger YAP activation through TRIP6 by increasing TRIP6 recruitment to cell-cell junctions, and TRIP6 binding to LATS1.

We also tested whether loss of tension across confluent (but not dense) monolayers of cells could trigger loss of LATS1-TRIP6 binding and colocalization at cell-cell junctions. Treatments that inhibit stress fibers such as type II myosin inhibition (Blebbistatin), Rho kinase inhibition (Y27632), or serum starvation are known to reduce tension at cell junctions (Yonemura et al., 2010). All of these treatments inhibited both LATS1-TRIP6 binding and cell-cell junction localization, as did complete elimination of F-actin using Latrunculin B (Figure 4A-C, S4A). These treatments (with the exception of Latrunculin B) did not obviously affect cell-cell adhesion and E-cadherin localization (Figure S4B). To reduce tension at cell junctions by blocking force transmission between cells, we disrupted cadherin complexes by treating cells with EGTA and found that this treatment also inhibited LATS1-TRIP6 binding and localization to cell-cell junctions (Figure 4A-C, S4A). Together these observations suggest that TRIP6 responds to tension at cell-cell junctions to regulate LATS1 and YAP activity.

**Figure 4.**
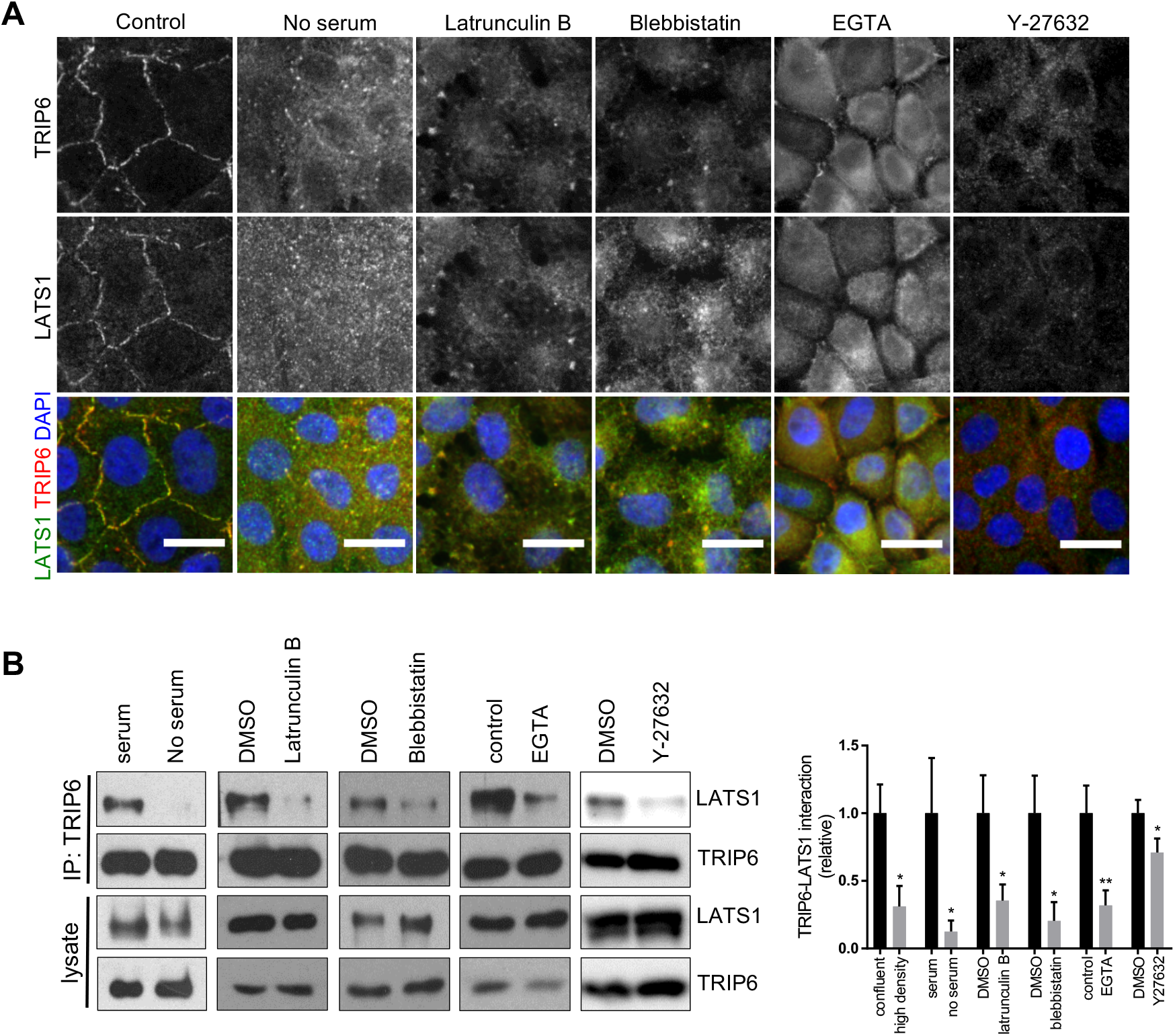
Perturbations of junctions and f-actin reduces TRIP6-LATS1 binding and localization to cell-cell junctions. (A) MCF10A cells were either not treated or treated separately with Latrunculin B, Blebbistatin, EGTA, serum starvation, and Y27632, and were stained for TRIP6 and LATS1. Merged images show LATS1 (green), TRIP6 (red) and DNA (blue). Scale bar=20µm. (B) Cells were treated as in (A), then lysed and anti-TRIP6 or control (IgG) antibodies were used to isolate immune complexes. Immune complexes and lysates were probed by western blotting for LATS1 and TRIP6. Quantification is shown. (Mean ± SD; n=3; *P<0.05, **P<0.01, T-test).

### TRIP6 is a part of the mechanosensory complex at adherens junctions

We next investigated whether TRIP6 could be part of a mechano-sensing complex at cadherin-catenin based adherens junctions (Huveneers and de Rooij, 2013). Previous studies showed that TRIP6 localizes to adherens junctions, and its association with the cadherin complex is dependent on engagement between the extra-cellular domains of cadherins on neighboring cells (Guo et al., 2014). How TRIP6 interacts with the cadherin complex is not known. Interestingly, two high throughput two-hybrid studies detected a binding interaction between TRIP6 and the adherens junction protein vinculin (Rual et al., 2005; Yu et al., 2011). Consistent with the high throughput studies, we detected vinculin in TRIP6 immune-complexes (Figure 5A). Vinculin and TRIP6 are known to respond to mechanical cues at focal adhesions (Bays and DeMali, 2017; Kuo et al., 2011; Schiller et al., 2011) and vinculin localization to adherens junctions depends on myosin contractile activity (Yonemura et al., 2010). As with TRIP6, we observed vinculin at focal adhesions in MCF10A cells at low density or at the edge of monolayers, but at densities used in this study (confluent but still proliferating), vinculin was concentrated at adherens junctions (Figure S5A). Thus, we infer that TRIP6 is primarily interacting with vinculin at adherens junctions under these conditions. At high cell density vinculin localization to adherens junctions was lost, but could be restored by stretch (Figure 5B-C), as observed for TRIP6 and LATS1. Vinculin localization to adherens junctions and binding to TRIP6 is reduced by treatments that disrupt tension (Figure 5B, 5D, S5B). In addition, when vinculin levels were reduced by siRNA, TRIP6-LATS1 binding and localization to cell-cell junctions were inhibited (Figure 5E-F; S5C-D). Depletion of vinculin also reduced YAP nuclear localization and activity (as judged by reduction in YAP target gene (CTGF and Cyr61) expression in MCF10A cells (Figure 5G-H). This same effect on YAP was observed using two different siRNAs (Figure S5E-F). Knockdown of vinculin in HEK293A cells also reduced YAP activity (S5G) and this reduction could be rescued by expression of vinculin (Figure S5G-H). Together these results show that tension stimulates vinculin recruitment to adherens junctions and binding to TRIP6. Vinculin promotes TRIP6-LATS1/2 binding and localization to adherens junctions to control YAP activity in response to changes in tension across tissues.

**Figure 5.**
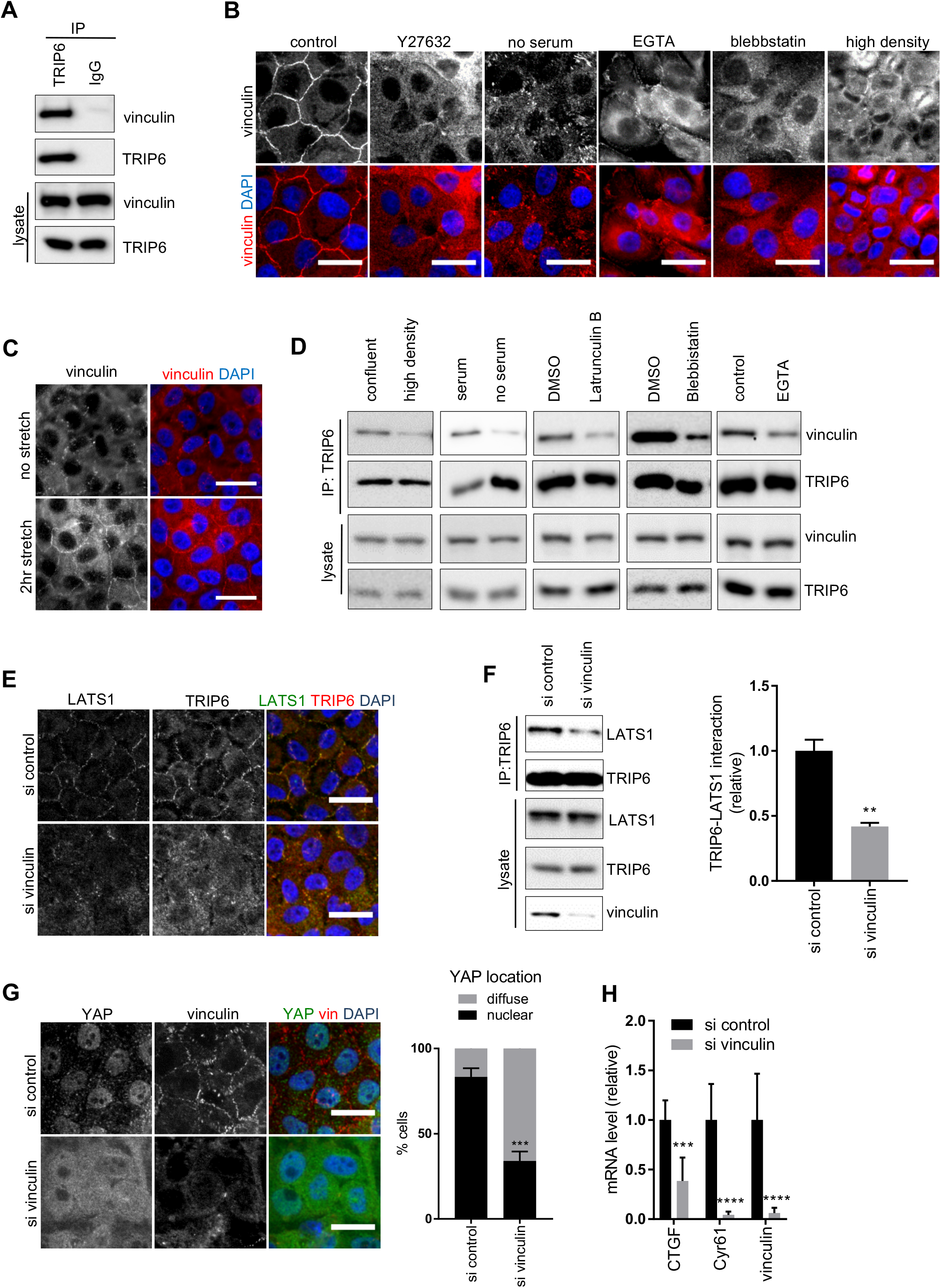
Vinculin regulates TRIP6-LATS1 interaction, localization, and YAP activity. (A) MCF10A cells were lysed and anti-TRIP6 or control (IgG) antibodies were used to isolate immune complexes. Immune complexes and lysates were probed by western blotting for vinculin and TRIP6. (B) MCF10A cells were either not treated (control) or treated separately by growth to high density, serum starvation, Latrunculin B, Blebbistatin, EGTA, or Y27632 treatment, and stained using anti-vinculin antibodies by immunofluorescence. Scale bar=20µm. (C) MCF10A cells grown at high density on PDMS membranes and were stretched (or not) at 17% elongation for 2 hours and fixed while under tension and stained for vinculin. Scale bar=20µm. (D) MCF10A cells were subjected to different treatments as described in Figure 4A and TRIP6 immune complexes and lysates were probed by western blotting for vinculin and TRIP6 (quantification is shown in Figure S5B). (E) Vinculin was knocked down using two different siRNAs or control siRNA in MCF10A cells and cells were stained for LATS1 and TRIP6. Merged images show LATS1 (green), TRIP6 (red) and DNA (blue). Scale bar=20µm. (F) Vinculin was knocked down as described in (E) and TRIP6 immune complexes and lysates were probed by western blotting for LATS1 and TRIP6 and the relative levels quantified (Mean ± SD; n=3; **P<0.01, T-test). (G) Vinculin was knocked down as described in (E) and cells were stained for YAP and TRIP6. Merged image shows YAP (green), TRIP6 (red), and DNA (blue). Quantification of at least 100 cells is shown (Mean ± SD; n=3; ***P<0.001, Fisher’s test). Scale bar=20µm. (H) Vinculin was knocked down as described in (E) and the levels of vinculin and YAP target gene expression was analyzed using RT-qPCR (Mean ± SD; n=3; ***P<0.001, ****P<0.0001, T-test).

## Discussion

This study provides new insight into the mechanism by which mechanical forces regulate cell growth and proliferation decisions via the Hippo signaling pathway. In particular, we report that the LIM domain protein TRIP6 functions as an intermediate between the LATS1/2 protein kinases, which transmit signals to YAP, and the mechano-responsive protein vinculin at the adherens junctions. Previous studies have shown that YAP activity can be stimulated by tension (Aragona et al., 2013; Benham-Pyle et al., 2015; Codelia et al., 2014; Rauskolb et al., 2014), however, the upstream signaling pathways remained uncertain. We found that tension stimulates TRIP6 binding to LATS1/2, and, once bound, TRIP6 inhibits LATS1/2 activity, reminiscent of work in *Drosophila* showing that the Ajuba LIM domain protein activates Yki (the YAP homolog) by inhibiting Warts (the LATS1/2 homolog) in response to tension (Rauskolb et al., 2014). Furthermore, we identified a specific molecular mechanism for how TRIP6 inhibits LATS1/2. We discovered that TRIP6 competes with MOB1 for binding to LATS1/2. MOB1 promotes LATS1/2 activation by scaffolding interactions between the LATS1/2 activating kinase MST1/2 and LATS1/2. The competition we observe between TRIP6 and MOB1 for binding to LATS1/2 is consistent with our other results showing TRIP6 interferes with MST1/2 phosphorylation of LATS1/2. This mechanism may be relevant for other LIM domain proteins that bind to a similar region of LAtS1/2 (Abe et al., 2006; Hirota et al., 2000), and could function in conjunction with other proposed mechanisms for how LIM domain proteins inhibit LATS1/2 (Jagannathan et al., 2016; Ma et al., 2016; Sun et al., 2015).

Our studies identified vinculin as acting upstream of TRIP6. Both TRIP6 and vinculin loss of function inhibits YAP activity, vinculin binds TRIP6, and is required to recruit TRIP6 and LATS1/2 to adherens junctions and promote their binding to each other. It is not clear how vinculin promotes TRIP6-LATS binding, but one possibility is that vinculin directly or indirectly causes a conformational change in TRIP6 to allow it to bind LATS1/2. Vinculin itself responds to mechanical tension at adherens junctions since both vinculin localization to adherens junctions and vinculin-TRIP6 binding is dependent on mechanical tension. Previous studies have shown that vinculin is recruited to adherens junctions by α-catenin, which responds directly to mechanical tension. Alpha-catenin binding to vinculin stabilizes its open conformation allowing it to bind actin and possibly other effectors like TRIP6 (Choi et al., 2012; Huveneers and de Rooij, 2013; Twiss et al., 2012; Yonemura et al., 2010). The open conformation of vinculin induced by α-catenin may stimulate vinculin binding to TRIP6. If this was the case then one would expect α-catenin and vinculin loss of function to have similar effects on YAP activity. However, previous studies showed that α-catenin loss of function stimulates YAP activity (Kim et al., 2011; Schlegelmilch et al., 2011; Silvis et al., 2011), in contrast to the increased YAP activity we observe when vinculin is knocked down. This apparent discrepancy could be resolved if adherens junctions and α-catenin had different functions in cells at lower density (higher tension) compared to cells at high density (low tension). It should be noted that the earlier studies showing α-catenin acting as an inhibitor of YAP were done at high cell density where tension would be low and vinculin and TRIP6 would not be at adherens junctions. At lower cell density, when cells are confluent but still proliferating (and presumably under more tension), α-catenin may recruit vinculin to adherens junctions to enhance YAP activity (via TRIP6 inhibition of LATS1/2) and drive cell proliferation. Thus, as cell density increases and tension decreases the vinculin-TRIP6 system turns off, and the YAP inhibitory function of adherens junctions could become dominant. It will be interesting in the future to determine how these two systems interact with each other to tune YAP regulation in response to changes in cell density and/or tension. In summary, we showed that TRIP6 acts as an intermediary connecting tension monitoring at adherens junctions to Hippo signaling, which has implications for how tension contributes to growth of organs and tissues during development, tissue repair during injury and to pathological conditions such as cancer.

## Author Contributions

S.D., S.M.C., and D.M. devised the experiments. S.D. performed most of the experiments with assistance from S.M.C., I.D., and H.C. S.M.C. made the TRIP6 knock out line in HEK293A cells and M.P. made LATS2 plasmids. K.B. and H.C. helped design the static stretch experiments. S.D., S.M.C., and D.M. wrote the manuscript.

## Acknowledgments

We thank Anthony Schmitt, Maria Fernandes, Elizabeth Luna, and Greenfield Sluder for antibodies, and Tom Fazzio for CRISPR plasmids. We thank Peter Pryciak for giving valuable feedback on the manuscript. This work was supported by the National Institutes of Health grant GM058406-18 to DM, and a UMMS-WPI seed grant to DM and KB.

## Supplemental Figures

**Figure S1.**

TRIP6 knockdown in MCF10A cells activates hippo signaling and TRIP6Δ knockout cells are rescued by FLAG-TRIP6 expression. (A) Lysates from MCF10A cells infected with control lentivirus (shEGFP) or lentivirus expressing shRNA against TRIP6 (shTRIP6) were analyzed by western blotting using the indicated antibodies, and the levels of LATS1 activating phosphorylation was quantified (Mean ± SD; n=3; **P<0.01, T-test). (B) MCF10A cells were infected with lentivirus carrying control shRNA (shEGFP), or two different shRNA against TRIP6 (shTRIP6-1, shTRIP6-4) and the levels of TRIP6 and YAP target gene expression was analyzed using RT-qPCR (Mean ± SD; n=3; **P<0.01, ***P<0.001, T-test). (C) MCF10A cells were infected with lentivirus carrying control shRNA (shEGFP), or a mix of two different shRNA against TRIP6 (shTRIP6-1 and shTRIP6-4) and were stained for YAP and TRIP6. Merged image shows YAP (green), TRIP6 (red), and DNA (blue). Quantification of YAP nuclear localization is shown (Mean ± SD; n=3; ****P<0.0001, Fisher’s test). Scale bar=20µm. (D) YAP, TRIP6 and tubulin levels were measured by western blotting in MCF10A cells infected with lentivirus carrying control shRNA (shEGFP), or shRNA against TRIP6 (shTRIP6-1) (Mean ± SD; n=3; **P<0.01, T-test). (E) Control (WT) and TRIP6Δ HEK293A cells were transfected with 200ng of control and FLAG-TRIP6 plasmids. (Note that 200ng of FLAG-TRIP6 plasmid restores approximate wild-type levels (see (F)) of TRIP6 expression). After 48 hours of transfection, cells were stained using anti-YAP and TRIP6 antibodies by immunofluorescence. Quantification of YAP nuclear localization is shown (Mean ± SD; n=3; ****P<0.0001, Fisher’s test). Scale bar=20µm. We compare TRIP6Δ cells to TRIP6Δ cells rescued by FLAG-TRIP6 plasmid (rescue) (F) Different amounts (50, 100, 150, 200ng) of FLAG-TRIP6 plasmid were transfected into HEK293A TRIP6Δ cells and TRIP6 levels in lysates were analyzed by western blotting using anti-TRIP6 antibodies and compared to those in control HEK293A (WT) cells. 200ng of FLAG-TRIP6 plasmid (marked with asterisk) was selected to perform the rescue experiment described in (E).

**Figure S2.**

Coomassie stained gel showing the amounts of BSA and highest amount of HIS-MOB1A used in Figure 2B.

**Figure S3.**

Regulation of TRIP6 and LATS1 localization and binding. (A) MCF10A cells were stained for TRIP6 and LATS1 and the image was acquired with a confocal microscope. Merged images show LATS1 (green), TRIP6 (red) and DNA (blue). Scale bar=20µm. The LATS1-TRIP6 colocalization was confirmed by line scan analysis of the pixel intensity in different fluorescent channels. (B) HEK293A cells (WT and TRIP6Δ) were grown and TRIP6 and LATS1 intracellular localization were determined by immunofluorescence using anti-TRIP6 and anti-LATS1 antibodies. HEK293A (WT) cells were also grown at high density and treated similarly. The localization of TRIP6 and LATS1 in HEK293A TRIP6Δ cells is also shown. Scale bar=20µm. (C) MCF10A cells were grown at low density and until confluency and stained for TRIP6 and FAK. Merged images show FAK (green), TRIP6 (red) and DNA (blue). (D) Quantification of LATS1 at cell-cell junctions in MCF10A depleted of TRIP6 (see Figure 3A). Intensity of LATS1 was measured at cell-cell junctions (Mean ± SD; n=3; **P<0.01, T-test). (E) Quantification of TRIP6 at cell-cell junctions in MCF10A depleted of LATS1 and LATS2 (see Figure 3B). Intensity of TRIP6 was measured at cell-cell junctions (Mean ± SD; n=3; n.s.≥0.05, T-test). (F) MCF10A cells grown at high density on PDMS membranes and were stretched (or not) at 17% elongation for 2 hours, RNA was isolated and YAP target gene expression was analyzed using RT-qPCR. (Mean ± SD; n=3; **P<0.01, ****P<0.0001, T-test).

**Figure S4.**

Regulation of TRIP6 and LATS1 localization in HEK293A cells and E-cadherin staining in MCF10A cells after various treatments. (A) HEK293A cells were either not treated (control) or treated separately by serum starvation (no serum), Latrunculin B, Blebbistatin, and EGTA, then TRIP6 and LATS1 intracellular localization were determined by immunofluorescence using anti-TRIP6 and anti-LATS1 antibodies. The localization of TRIP6 and LATS1 in HEK293A TRIP6Δ cells is also shown. Scale bar=20µm. (B) MCF10A cells were either not treated (control) or treated separately by growth to high density, serum starvation, Latrunculin B, Blebbistatin, or Y27632 treatment, and stained using anti-E-cadherin antibodies by immunofluorescence. Scale bar=20µm.

**Figure S5.**

FAK and vinculin co-staining in MCF10A cells, quantification of TRIP6-vinculin binding and vinculin knockdown efficacy, LATS1 and vinculin co-staining in MCF10A cells, effect of vinculin knockdown by single siRNAs on YAP activity and localization and rescue of siRNA knockdown are shown. (A) MCF10A cells were grown at low density and until confluency and stained using anti-FAK and anti-vinculin antibodies by immunofluorescence. Scale bar=20µm. (B) Quantification of the TRlp6-vinculin binding from Figure 5D is shown. The relative TRIP6-vinculin binding was determined for MCF10A cells under different conditions compared to controls (confluent/high density, +/-serum, +/-Latrunculin B, +/-Blebbistatin, or +/-EGTA). (Mean ± SD; n=3; *P<0.05, **P<0.01, ***P<0.001, T-test) (C) Vinculin was knocked down using two different stealth siRNAs in MCF10A cells. MCF10A control cells were treated with control siRNA. Lysates were probed by western blotting for vinculin and tubulin. The levels of vinculin were determined. (Mean ± SD; n=3; ***P<0.001, T-test) (D) Vinculin was knocked down as described in (C) and cells were stained using vinculin and LATS1 antibody. Merged images show LATS1 (green), vinculin (red) and DNA (blue). Scale bar=20µm. (E) Vinculin was knocked down by single stealth siRNAs (7662 and 1260) in MCF10A cells and stained using vinculin and YAP antibody. Merged images show YAP (green), vinculin (red) and DNA (blue). Quantification of YAP nuclear localization is shown (Mean ± SD; n=3; **P<0.01, ***P<0.001, Fisher’s test). Scale bar=20µm. (F) Vinculin was knocked down as described in (E) and the levels of vinculin, and YAP target gene expression was analyzed using RT-qPCR. (Mean ± SD; n=3; **P<0.01, ***P<0.001, T-test). (G) Vinculin was knocked down as described in 5E in HEK293A cells. Control and vinculin depleted HEK293A cells were transfected with 150ng of control and EGFP-vinculin plasmids. (Note that 150ng of EGFP-vinculin plasmid restores approximate wild-type levels (see (H)) of vinculin expression). After 48 hours of transfection, the levels of vinculin, and YAP target gene expression was analyzed using RT-qPCR. We compare vinculin depleted cells to vinculin depleted cells rescued by EGFP-vinculin (rescue). (Mean ± SD; n=3; *P<0.05, **P<0.01, ****P<0.0001, T-test). (H) Vinculin was knocked down as described in 5E in HEK293A cells. Different amounts (50, 150, 250, 350ng) of EGFP-vinculin plasmid were transfected into the vinculin depleted cells and vinculin levels in lysates were analyzed by western blotting using anti-vinculin antibody and compared to those in control HEK293A cells. 150ng of EGFP-vinculin plasmid (marked with asterisk) was selected to perform the rescue experiment.

## Supplemental Materials and Methods

### KEY RESOURCES TABLE

**Table S1.**
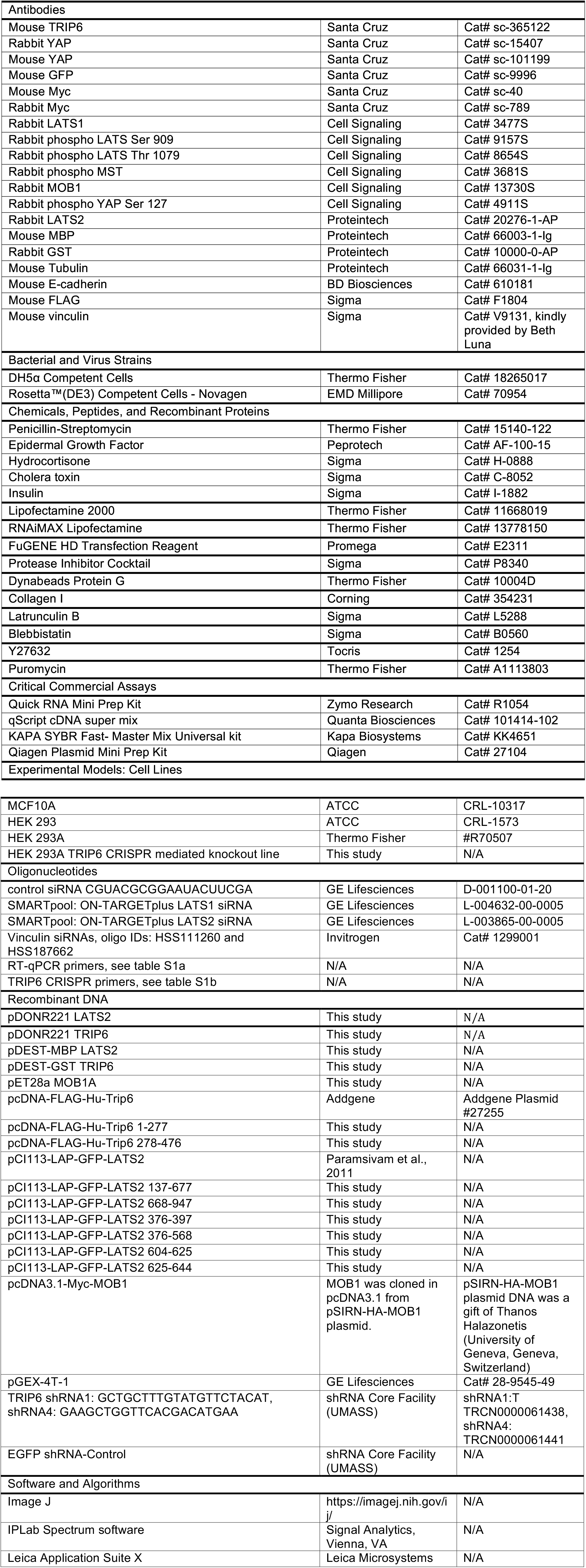
Table S1 consists of a list of qPCR primers and primers for CRISPR mediated TRIP6 knockout in HEK293A cells.

### CONTACT FOR REAGENT AND RESOURCE SHARING

Further information and requests for reagents may be directed to and will be fulfilled by the Lead Contact, Dannel McCollum.

### EXPERIMENTAL MODEL AND SUBJECT DETAILS

#### Cell Lines

Human Embryonic Kidney (HEK293, HEK293A) cell lines were grown in Dulbecco’s modified Eagle medium (DMEM, GIBCO) supplemented with 10% (v/v) fetal bovine serum (FBS, GIBCO) and 1% (v/v) penicillin/streptomycin (Invitrogen). Human mammary epithelial cell line MCF10A was cultured in DMEM/F12 (1:1) media supplemented with 5% (v/v) fetal horse serum (GIBCO), 20ng/ml Epidermal Growth Factor (Peprotech), 0.5 mg/ml Hydrocortisone (Sigma), 100 ng/ml Cholera toxin (Sigma), 10µg/ml Insulin (Sigma) and 1% (v/v) penicillin/streptomycin (Invitrogen). Cell lines were cultured in a humidified incubator at 37°C with 5% CO2.

#### Expression Plasmids and shRNAs

Flag-TRIP6 plasmid is obtained from Addgene (Plasmid #27255). shRNAs for TRIP6 (shRNA1: TRCN0000061438, shRNA4: TRCN0000061441) and control shRNA (shEGFP) were obtained from the UMass RNAi core facility.

### METHOD DETAILS

#### CRISPR mediated deletion of TRIP6 in 293A cell line

The target sequence to knock-out the TRIP6 gene was selected with the web tool developed by the Zhang lab (http://crispr.mit.edu/). Oligos complementary to the target sequence with appropriate overhangs (CACCGGCGATCCCCCGCGGCACCC and AACGGGTGCCGCGGGGGATCGCC) were annealed and cloned into a variant of the px330 plasmid with puromycin resistance (Hainer et al., 2015). HEK293A and HEK293 cells were transfected with Lipofectamine 2000 (Invitrogen) following the manufacturer directions and using 500 ng of plasmid per well of a 12-well plate. The next day, cells were placed under selection with 2 µg/ml of Puromycin (GIBCO) for 48 hours. Puromycin resistant cells were then heavily diluted and plated on 10 cm plates for colony isolation. Colonies were picked 7-10 days later by depleting the media from the plate and using a P20 pipette loaded with 1 µl of media to dislodge the colony from the plate by rapid back and forth movements. Clonal lines were then expanded and the expression of TRIP6 was determined by Western blot. Clonal lines that lacked expression of TRIP6 were again expanded from single cells by dilution followed by colony isolation and tested by Western blot to ensure that they were true clonal lines. At least two independent clonal lines were kept for further analysis. HEK293A cells were maintained at low densities as much as possible to prevent morphological changes associated with cell crowding.

#### Cell starvation and drug treatments

MCF10A and HEK293A cells were starved overnight in DMEM/F12 (1:1) and DMEM respectively supplemented with 1% of penicillin and streptomycin (Invitrogen) before adding complete cell culture medium described above for 1 hour. Latrunculin B was used at 1 µM for 1 hour. Blebbistatin was used at 25 µM for 2 hours on MCF10A and 1 hour on 293A. Both MCF10A and 293A cell lines were treated with 0.5mM EGTA for 30 minutes. MCF10A cells were treated with 50µM of Y27632 for 1 hour.

#### Stretching experiments

MCF10A cells were cultured on collagen I coated bioflex plates (Flexcell International Corporation (# BF-3001C)) at high density before stretching them with a Flexcell FX-4000 machine (Flexcell International, Burlington, NC) using 22mm diameter posts under maximum vacuum pressure, resulting in a 17% equibiaxial stretch for 2 hours in a humidified incubator at 37°C with 5% CO2. Cells were either lysed for RNA preparation (see RT-qPCR), protein preparation (Cell lysis, Immunoprecipitation and Western Blotting) or fixed while under stretch with 3.7% PFA for 10 minutes before performing immunofluorescence. Control plates were not stretched. For stretched plates, the stretch level was validated by measuring deformation before and after stretch using multiple fiduciary markers.

#### Immunofluorescence

HEK 293A and MCF10A cells were cultured on coverslips and fixed with 3.7% paraformaldehyde in PBS for 10 minutes, permeabilized in 0.5% Triton-X in UB (UB; 150 mM NaCl, 50 mM Tris pH 7.6, 0.01% NaN3) for 3 minutes at 37°C, then blocked with 10% BSA in UB for 30 minutes at 37°C. Cells were then incubated for 1 hour at 37°C with appropriate primary antibodies, washed three times in UB and incubated with Alexa Fluor-conjugated secondary antibodies (Molecular probes) for 1 hour at 37°C. After three washes in UB, coverslips were mounted on glass slides using Prolong Gold Antifade reagent with DAPI (Invitrogen) and left at 4°C overnight. The next day slides were viewed using fluorescent microscopy (Nikon Eclipse E600) and images were acquired using a cooled charge-coupled device camera (ORCA-ER; Hamamatsu, Bridgewater, NJ). The confocal image was acquired using a Leica SP5 AOBS second generation laser scanning confocal microscope. Image processing and analysis were carried out with IPLab Spectrum software (Signal Analytics, Vienna, VA) and ImageJ software (Schneider et al., 2012).

#### Cell lysis, Immunoprecipitation and Western Blotting

HEK293 and HEK293A cells were transfected using Lipofectamine 2000 (Invitrogen) according to the manufacturer’s protocol. For the rescue assay, HEK293A (WT) and HEK293A TRIP6Δ knockout cells were transfected with empty plasmid and increasing amounts of FLAG-TRIP6 plasmid (50ng-200ng) respectively, using FuGENE^®^ HD Transfection Reagent (#E2311, Promega) according to manufacturer’s protocol. 200ng of the plasmid was used for the final rescue experiment. Cells were collected after 48 hours and lysed with lysis buffer (10% Glycerol (Invitrogen), 20mM Tris-HCl-pH 7.0, 137mM NaCl, 2mM EDTA, 1% NP-40 (Invitrogen), 1mM PMSF(Sigma), 1mM Na3VO4 (Sigma) and 1x mammalian protease inhibitor cocktail (Sigma)). MCF10A cells were additionally passed through a 26G½ needle. Cells were then incubated for 10 minutes at 4°C and lysates were cleared by centrifugation at 10,000 rpm for 10 min at 4°C. For immunoprecipitation, Dynabeads (Invitrogen) were used according to the manufacturer’s protocol.

#### siRNA/shRNA transfection

Knockdowns in MCF10A cells were performed using 30 nM of control siRNA (fire fly luciferase) or SMARTpool siRNA from Dharmacon (for LATS1 and 2) or stealth siRNA from Thermo-Fisher (for vinculin). RNAiMAX Lipofectamine (Invitrogen) was used according to the manufacturer’s protocol. After 48 hours cells were either used for western blotting or fixed for immunofluorescence. Stable knockdowns in MCF10A cells were done using lentiviral infection of shRNA and cells were selected with puromycin for 3 days. Experiments were performed immediately after puromycin selection. Viral supernatants were generated by the shRNA Core Facility (UMASS) to target TRIP6.

#### RT-qPCR

RNA was prepared using Quick-RNA MiniPrep kit (Zymo Research) according to the manufacturer’s protocol. cDNA was prepared using qScript cDNA super mix (Quanta Biosciences, Inc) according to the manufacturer’s protocol. RT-qPCR was performed using KApA SYBR Fast-Master Mix Universal kit (Kapa Biosystems). Target mRNA levels were measured relative to GAPDH mRNA levels.

#### Recombinant protein expression and ***in vitro*** competition assays

TRIP6 and LATS2 were cloned in pDEST-GST and pDEST-MBP respectively (provided by Dr. Marian Walhout’s lab) using Gateway (ThermoFisher Scientific) directions. MOB1A was cloned in pET28a through standard cloning. GST-TRIP6, MOB1A-6xHis, and MBP-LaTS2 plasmids were transformed into BL21 DE3 cells and recombinant protein expression was induced with 1mM IPTG for 4 hours and 30 minutes at 25° C. Bacterial pellets were resuspended in lysis buffer (1.8 mM KH_2_PO_4_,10 mM Na_2_HPO_4_, 150 mM NaCl, 10 mM β-mercapto-ethanol, 0.05% Triton X-100, 1 mg/ml of lysozyme, 5 µg/ml of nuclease, and 1mM PMSF) and incubated for 30 minutes at 4° C. Cells were lysed on ice with 6 rounds of 10 sonications each using a VWR Sonifier 450 fitted with a microtip set to an output of 2 and a duty cycle of 80. Lysates were cleared by centrifugation at 21,000g for 10 minutes at 4° C. GST-TRIP6 was purified using Glutathione beads (GE) and eluted with 20 mM glutathione for 30 minutes at 4° C in elution buffer (1.8 mM KH_2_PO_4,10_ mM Na_2_HPO_4_, 150 mM NaCl, 0.05% Triton X-100, and 1mM PMSF). MOB1A-6His was purified using Ni-IDA beads (Biotool) and eluted with 300 mM Imidazole for 30 minutes at 4° C in elution buffer. MBP-LATS2 was pulled down with magnetic maltose beads (NEB). For the in vitro competition assay, GST-TRIP6, MOB1A-6xHis, and control proteins were mixed as indicated in Figure 3B and then adjusted to a volume of 60µl using elution buffer. A constant amount of GST-TRIP6 (approximately 0.7 µg) was used in each sample, and either 1, 4, or 10-fold molar ratios of MOB1A-6xHis were added as competitor. The different protein solutions were then added with 20 µl of 10 mM Tris-HCl, pH 7.4 (to ensure an equal pH) to bead bound MBP-LATS2, and incubated for 20 minutes at room temperature with mixing. MBP-LATS2 bound beads were separated using a magnetic stand, washed 3 times in elution buffer, and boiled in SDS-PAGE sample buffer. Protein samples were then subjected to SDS-PAGE and Western blotting with the specified antibodies.

### QUANTIFICATION AND STATISTICAL ANALYSIS

Data are presented as Mean ± SD. Each experiment was done in triplicates. Students t-test (*P ≤ 0.05, ** P ≤ 0.01, *** P ≤ 0.001, **** P ≤ 0.0001) were performed using Prism version 7.00 for Windows (GraphPad Software, La Jolla California USA, www.graphpad.com). For YAP localization studies, we counted 100 cells each from three different experiments and used Fisher’s test (* P ≤ 0.05, ** P < 0.01, *** P ≤ 0.001, **** P ≤ 0.0001) using GraphPad Quickcalcs (http://graphpad.com/quickcalcs/contingency1/) to calculate the significance. For western blots, we performed background substraction and densitometric analysis of respective bands using Image J (Schneider et. al., 2012) and normalized to loading control (either actin or tubulin).

## REFERENCES

Abe Y., M. Ohsugi, K. Haraguchi, J. Fujimoto, and T. Yamamoto. 2006. LATS2-Ajuba complex regulates gamma-tubulin recruitment to centrosomes and spindle organization during mitosis. FEBS Lett. 580:782–788.

Aragona M, T. Panciera, A. Manfrin, S. Giulitti, F. Michielin, N. Elvassore, S. Dupont, and S. Piccolo. 2013. A Mechanical Checkpoint Controls Multicellular Growth through YAP/TAZ Regulation by Actin-Processing Factors. Cell. 154:1047–1059.

Bays J.L., and K.A. DeMali. 2017. Vinculin in cell-cell and cell-matrix adhesions. Cell Mol Life Sci.

Benham-Pyle B.W., B.L. Pruitt, and W.J. Nelson. 2015. Cell adhesion. Mechanical strain induces E-cadherin-dependent Yap1 and beta-catenin activation to drive cell cycle entry. Science. 348:1024–1027.

Chastre E, M. Abdessamad, A. Kruglov, E. Bruyneel, M. Bracke, Y. Di Gioia, M.C. Beckerle, F. van Roy, and L. Kotelevets. 2009. TRIP6, a novel molecular partner of the MaGI-1 scaffolding molecule, promotes invasiveness. FASEB J. 23:916–928.

Codelia V.A., G. Sun, and K.D. Irvine. 2014. Regulation of YAP by mechanical strain through Jnk and Hippo signaling. Curr Biol. 24:2012–2017.

Das Thakur, M., Y. Feng, R. Jagannathan, M.J. Seppa, J.B. Skeath, and G.D. Longmore. 2010. Ajuba LIM proteins are negative regulators of the Hippo signaling pathway. Curr Biol. 20:657–662.

Fei J, J. Li, S. Shen, and W. Zhou. 2013. Characterization of TRIP6-dependent nasopharyngeal cancer cell migration. Tumour Biol. 34:2329–2335.

Grunewald T.G., S. Willier, D. Janik, R. Unland, C. Reiss, O. Prazeres da Costa, T. Buch, U. Dirksen, G.H. Richter, F. Neff, S. Burdach, and E. Butt. 2013. The Zyxin-related protein thyroid receptor interacting protein 6 (TRIP6) is overexpressed in Ewing’s sarcoma and promotes migration, invasion and cell growth. Biol Cell. 105:535–547.

Guo Z., L.J. Neilson, H. Zhong, P.S. Murray, S. Zanivan, and R. Zaidel-Bar. 2014. E-cadherin interactome complexity and robustness resolved by quantitative proteomics. Sci Signal. 7:rs7.

Halder G, S. Dupont, and S. Piccolo. 2012. Transduction of mechanical and cytoskeletal cues by YAP and TAZ. Nat Rev Mol Cell Biol. 13:591–600.

Heller E., and E. Fuchs. 2015. Tissue patterning and cellular mechanics. J Cell Biol. 211:219–231.

Hirota T, T. Morisaki, Y. Nishiyama, T. Marumoto, K. Tada, T. Hara, N. Masuko, M. Inagaki, K. Hatakeyama, and H. Saya. 2000. Zyxin, a regulator of actin filament assembly, targets the mitotic apparatus by interacting with hwarts/LATS1 tumor suppressor. J Cell Biol. 149:1073–1086.

Huang S., and D.E. Ingber. 1999. The structural and mechanical complexity of cell-growth control. Nat Cell Biol. 1:E131-138.

Huveneers S., and J. de Rooij. 2013. Mechanosensitive systems at the cadherin-F-actin interface. J Cell Sci. 126:403–413.

Jagannathan R., G.V. Schimizzi, K. Zhang, A.J. Loza, N. Yabuta, H. Nojima, and G. D. Longmore. 2016. AJUBA LIM proteins limit Hippo activity in proliferating cells by sequestering the Hippo core kinase complex in the cytosol. Mol Cell Biol.

Kuo J.C., X. Han, C.T. Hsiao, J.R. Yates, 3rd, and C.M. Waterman. 2011. Analysis of the myosin-II-responsive focal adhesion proteome reveals a role for beta-Pix in negative regulation of focal adhesion maturation. Nat Cell Biol. 13:383–393.

Liu C.Y., Z.Y. Zha, X. Zhou, H. Zhang, W. Huang, D. Zhao, T. Li, S.W. Chan, C.J. Lim, W. Hong, S. Zhao, Y. Xiong, Q.Y. Lei, and K.L. Guan. 2010. The hippo tumor pathway promotes TAZ degradation by phosphorylating a phosphodegron and recruiting the SCF{beta}-TrCP E3 ligase. J Biol Chem. 285:37159–37169.

Mammoto T, A. Mammoto, and D.E. Ingber. 2013. Mechanobiology and developmental control. Annu Rev Cell Dev Biol. 29:27–61.

Meng Z, T. Moroishi, and K.L. Guan. 2016. Mechanisms of Hippo pathway regulation. Genes Dev. 30:1–17.

Meng Z, T. Moroishi, V. Mottier-Pavie, S.W. Plouffe, C.G. Hansen, A.W. Hong, H. W. Park, J.S. Mo, W. Lu, S. Lu, F. Flores, F.X. Yu, G. Halder, and K.L. Guan. 2015. MAP4K family kinases act in parallel to MST1/2 to activate LATS1/2 in the Hippo pathway. Nat Commun. 6:8357.

Mui K.L., C.S. Chen, and R.K. Assoian. 2016. The mechanical regulation of integrin-cadherin crosstalk organizes cells, signaling and forces. J Cell Sci. 129:1093–1100.

Ni L, Y. Zheng, M. Hara, D. Pan, and X. Luo. 2015. Structural basis for Mob1-dependent activation of the core Mst-Lats kinase cascade in Hippo signaling. Genes Dev. 29:1416–1431.

Paramasivam M, A. Sarkeshik, J.R. Yates, 3rd, M.J. Fernandes, and D. McCollum. 2011. Angiomotin family proteins are novel activators of the LATS2 kinase tumor suppressor. Mol Biol Cell. 22:3725–3733.

Rauskolb C, G. Pan, B.V. Reddy, H. Oh, and K.D. Irvine. 2011. Zyxin links fat signaling to the hippo pathway. PLoS Biol. 9:e1000624.

Rauskolb C, S. Sun, G. Sun, Y. Pan, and K.D. Irvine. 2014. Cytoskeletal tension inhibits Hippo signaling through an Ajuba-Warts complex. Cell. 158:143156.

Reddy B.V., and K.D. Irvine. 2013. Regulation of Hippo signaling by EGFR-MAPK signaling through Ajuba family proteins. Dev Cell. 24:459–471.

Rual J.F., K. Venkatesan, T. Hao, T. Hirozane-Kishikawa, A. Dricot, N. Li, G.F. Berriz, F.D. Gibbons, M. Dreze, N. Ayivi-Guedehoussou, N. Klitgord, C. Simon, M. Boxem, S. Milstein, J. Rosenberg, D.S. Goldberg, L.V. Zhang, S.L. Wong, G. Franklin, S. Li, J.S. Albala, J. Lim, C. Fraughton, E. Llamosas, S. Cevik, C. Bex, P. Lamesch, R.S. Sikorski, J. Vandenhaute, H.Y. Zoghbi, A. Smolyar, S. Bosak, R. Sequerra, L. Doucette-Stamm, M.E. Cusick, D.E. Hill, F.P. Roth, and M. Vidal. 2005. Towards a proteome-scale map of the human protein-protein interaction network. Nature. 437:1173–1178.

Schiller H.B., C.C. Friedel, C. Boulegue, and R. Fassler. 2011. Quantitative proteomics of the integrin adhesome show a myosin Il-dependent recruitment of LIM domain proteins. EMBO Rep. 12:259–266.

Sun S., and K.D. Irvine. 2016. Cellular Organization and Cytoskeletal Regulation of the Hippo Signaling Network. Trends Cell Biol. 26:694–704.

Vogel V., and M. Sheetz. 2006. Local force and geometry sensing regulate cell functions. Nat Rev Mol Cell Biol. 7:265–275.

Wang Y., J.E. Dooher, M. Koedood Zhao, and T.D. Gilmore. 1999. Characterization of mouse Trip6: a putative intracellular signaling protein. Gene. 234:403–409.

Yonemura S, Y. Wada, T. Watanabe, A. Nagafuchi, and M. Shibata. 2010. alpha-Catenin as a tension transducer that induces adherens junction development. Nat Cell Biol. 12:533–542.

Yu F.X., B. Zhao, and K.L. Guan. 2015. Hippo Pathway in Organ Size Control, Tissue Homeostasis, and Cancer. Cell. 163:811–828.

Yu H, L. Tardivo, S. Tam, E. Weiner, F. Gebreab, C. Fan, N. Svrzikapa, T. Hirozane-Kishikawa, E. Rietman, X. Yang, J. Sahalie, K. Salehi-Ashtiani, T. Hao, M.E. Cusick, D.E. Hill, F.P. Roth, P. Braun, and M. Vidal. 2011. Next-generation sequencing to generate interactome datasets. Nat Methods. 8:478–480.

Zhang J., G.A. Smolen, and D.A. Haber. 2008. Negative regulation of YAP by LATS1 underscores evolutionary conservation of the Drosophila Hippo pathway. Cancer Res. 68:2789–2794.

Zhao B, L. Li, K. Tumaneng, C.Y. Wang, and K.L. Guan. 2010. A coordinated phosphorylation by Lats and CK1 regulates YAP stability through SCF(beta-TRCP). Genes Dev. 24:72–85.

Zhao M.K., Y. Wang, K. Murphy, J. Yi, M.C. Beckerle, and T.D. Gilmore. 1999. LIM domain-containing protein trip6 can act as a coactivator for the v-Rel transcription factor. Gene Expr. 8:207–217.

